# Learning dynamically regulates stimulus discrimination of ventral striatal D1 receptor expressing neurons

**DOI:** 10.1101/2025.11.21.689812

**Authors:** Tierney B. Daw, Jiayi Cao, Ruoxian Li, Sotiris C. Masmanidis

**Affiliations:** Molecular, Cellular, and Integrative Physiology Graduate Program, University of California, Los Angeles, CA, USA; Department of Neurobiology, University of California, Los Angeles, CA, USA; Department of Bioengineering, University of California, Los Angeles, CA, USA; California Nanosystems Institute, University of California, Los Angeles, CA, USA

## Abstract

Animals encounter a barrage of sensory stimuli, but only a subset of these are associated with appetitive outcomes, highlighting the importance of neural mechanisms for learning to distinguish reward-paired from unpaired cues. The ventral striatum plays a critical role in both reinforcement learning and stimulus discrimination, but the effect of learning on the selectivity of different cell types remains unclear. Here we examined ventral striatal D1 and D2 medium spiny neuron (MSN) firing properties as mice learned to distinguish between reward-paired and unpaired cues. As learning progressed within a single session, D1 MSN selectivity increased linearly with behavioral selectivity, while D2 MSNs exhibited only modest, behaviorally uncorrelated changes in activity. Altered D1 MSN selectivity was primarily attributed to attenuated excitatory responses to the unrewarded cue, and increasing D1 MSN activity during the unrewarded cue impaired behavioral selectivity. Together, these findings reveal significantly more dynamic contributions of D1 MSNs to stimulus discrimination learning.

---

From foraging for berries to choosing an ice cream flavor, behavior is often guided by past associations of specific sensory cues with appetitive experiences. The stakes can be exceedingly high – picking the wrong ice cream is disappointing, but eating the wrong berry may be deadly. As a result, many species have evolved neural systems for distinguishing cues that predict rewarded from unrewarded outcomes. A large body of work has shown that the ventral striatum plays a major role in this process^1–5^. Importantly, the ventral striatum is implicated in both reinforcement learning and stimulus discrimination, functions essential for acquiring and expressing accurate reward predictions. For example, ventral striatal neurons represent information related to rewards and associated predictive cues in an experience-dependent manner^6–8^. In stimulus discrimination tasks, ventral striatal neurons selectively respond to cues predictive of appetitive, aversive, or neutral outcomes, and these responses evolve during learning^9–11^. This brain area has also been shown to suppress unproductive or impulsive actions, such as movement in response to an unrewarded cue^12–14^. Collectively, these studies demonstrate the critical role of ventral striatum in learning to appropriately respond or withhold responding to specific predictive stimuli. However, the contribution of different ventral striatal cell types to stimulus discrimination learning is not well understood.

The striatum predominantly consists of GABAergic medium spiny projection neurons (MSNs). MSNs are canonically divided into two genetically and functionally non-overlapping subtypes that express D1 or D2 dopamine receptors (some overlap does exist^15^, but it represents a relatively small population of cells that are beyond the scope of the present study). D1 and D2 MSNs form distinct circuit pathways that, when activated, are capable of antagonistically regulating movement and reward seeking behaviors^16–20^. However, a clear understanding of how these distinct cell types evolve during learning has been elusive. This is potentially because, despite significant progress in characterizing D1 and D2 MSN dynamics in diverse behaviors^21–26^, efforts examining activity changes as a function of reward-guided learning have been more limited. Among the handful of such studies that exist, three reported similar trends between D1 and D2 MSNs during the acquisition of reward-guided tasks^27–29^, while a fourth concluded that D2, but not D1, MSNs scale their responses to reward-paired cues with learning^30^. Crucially, previous experiments were not designed to examine how learning shapes the stimulus discrimination properties of D1 and D2 MSNs.

Based on the lack of firm evidence, several conceptual models could explain how D1 and D2 MSN activity changes as animals learn to distinguish between reward-paired and unpaired cues (**Fig. 1a**). The models focus on neural selectivity, a measure of how effectively cells discriminate between the two predictive stimuli. According to the classical view of opponent behavioral functions^16,17,31^, one set of predictions is that D1 MSN selectivity surpasses that of D2 MSNs as learning unfolds (models 1-3). Simple variants of these models, whereby D2 surpasses D1 MSN selectivity, were not shown but may be predicted by some literature^30^. There is also substantial evidence that would favor a balanced change in D1 and D2 MSN selectivity (model 4)^21,27,28,32^. Finally, if these circuits have an innate or “hardwired” ability to discriminate different stimuli, we may expect to find no learning-dependent modifications to selectivity (model 5)^11^. Thus, while past work can help guide our predictions, it is far from clear how D1 and D2 MSN dynamics are modified during stimulus discrimination learning.

**Fig. 1.**
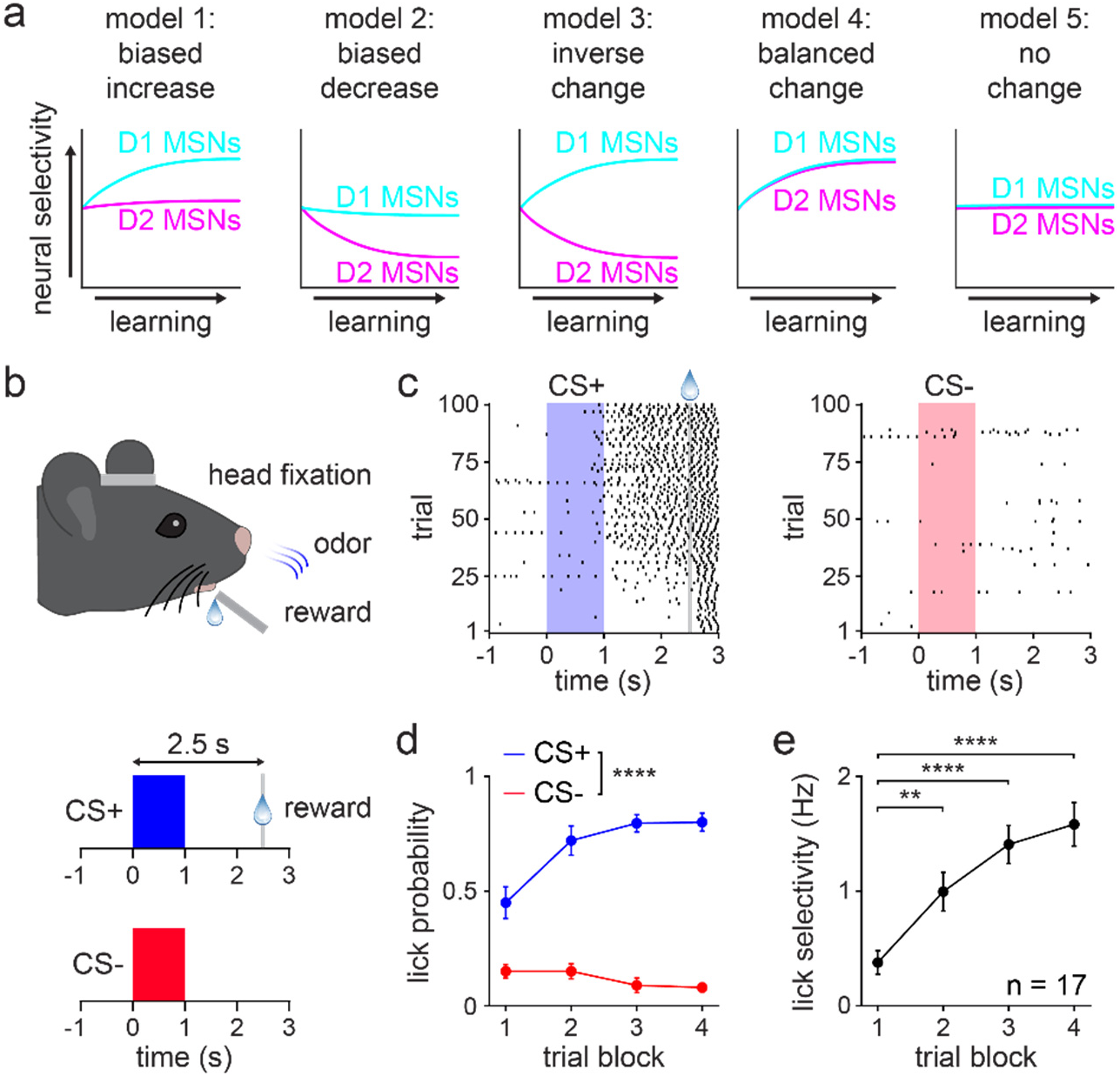
Mice learn to discriminate predictive cues in a single training session. **a.** Conceptual models illustrating a variety of possible learning-associated changes in striatal MSN activity that could underly the learning of stimulus discrimination. Variations of these models are also possible (i.e., a version of model 1 with D2 surpassing D1 MSN selectivity). **b.** Head-fixed behavioral setup for the classically conditioned cue discrimination task (top) and schematic of the task (bottom). CS+ was paired with reward on 100% of trials, and CS- was paired with reward on 0% of trials. **c.** Lick raster from one mouse on CS+ (left) and CS- (right) trials. **d.** CS+ and CS- lick probability across four 25-trial blocks (*n* = 17 mice; two-way RM ANOVA; block effect: *F*_3,96_ = 9, *P* < 0.0001; cue effect: *F*_1,32_ = 156, *P* < 0.0001; multiple comparisons for CS+ versus CS- on blocks 1-4 (*P* < 0.0001)). **e.** Lick selectivity versus trial block (one-way RM ANOVA; block effect: *F*_3,48_ = 22, *P* < 0.0001; multiple comparisons for blocks 1 versus 2 (*P* = 0.0011), 3 (*P* < 0.0001), and 4 (*P* < 0.0001)). d-e depict mean ± SEM. ** *P* < 0.01, **** *P* < 0.0001. See also Extended Data Fig. 1.

Here, we investigated whether any of these models are consistent with new experimental observations. Mice were trained on a classically conditioned stimulus discrimination task in which two randomly interleaved olfactory cues were consistently followed by either a reward or no outcome. During learning, animals increased behavioral selectivity, characterized by elevated anticipatory licking to the reward-paired cue and reduced licking to the unpaired cue. Discriminative behavior was acquired within a single training session, enabling us to track how learning impacted the activity of optogenetically identified D1 and D2 MSNs in the ventral striatum. Learning was primarily associated with increased D1 MSN cue selectivity, which was correlated with behavioral selectivity. By contrast, D2 MSNs displayed weaker changes in selectivity that were insignificantly correlated with behavior. These results are most consistent with the first conceptual model. Surprisingly, the increase in D1 MSN selectivity was mainly attributed to diminished modulation to the unrewarded cue rather than elevated modulation to the rewarded cue. This suggests that, as animals gain experience, stimulus discrimination is primarily enhanced by attenuating D1 MSN responses that are not associated with reward. Indeed, reversing this attenuation, by activating D1 MSNs specifically on unrewarded trials, impaired behavioral discrimination by altering licking to both types of cues in an opponent manner. Taken together, these findings indicate a significantly more dynamic role for ventral striatal D1 MSNs in learning to discriminate between reward-paired and unpaired stimuli.

## RESULTS

### Mice learn a stimulus discrimination task in a single session

Using a head-fixed classical conditioning paradigm, D1-Cre and A2a-Cre mice were trained to discriminate between two olfactory conditioned stimuli (**Fig. 1b**)^33^. One conditioned stimulus (CS+) was consistently followed by sweetened milk reward after a 2.5 s delay, while the other conditioned stimulus (CS-) was never paired with reward. The training session consisted of 100 trials of each cue presented in randomly interleaved order. Within a single training session, mice learned to lick in anticipation of reward following the CS+ and mostly suppressed licking following the CS- (**Fig. 1c**). To assess discriminative behavior across learning, the training session was parsed into four blocks of 25 trials of each cue. Across the four blocks, the anticipatory licking probability increased for CS+ trials and declined for CS- trials (**Fig. 1d**). Thus, the licking selectivity, calculated as the difference between CS+ and CS- evoked anticipatory lick rates, increased significantly as learning unfolded (**Fig. 1e**). These behavioral results combined data from D1-Cre and A2a-Cre mice after confirming that these groups did not exhibit significant differences in licking performance (**Extended Data Fig. 1a-c**). To verify that CS+ and CS- evoked behavior showed similar values early in learning, lick probability and selectivity were also quantified using 5-trial blocks across the first 25 trials of the training session. As expected, there was no significant difference between CS+ and CS- lick probabilities in the first 5-trial block (**Extended Data Fig. 1d**); likewise, the lick selectivity started near a value of zero (**Extended Data Fig. 1e**). These results confirmed that differences between CS+ and CS- evoked behavior were associated with learning, as opposed to being driven by an innate response to the olfactory stimuli. Moreover, mice successfully acquired the task in a single training session, enabling us to study the neurophysiological underpinnings of discriminative learning.

### Electrophysiological recordings of ventral striatal D1 and D2 MSNs

To track neural activity across learning, the same groups of mice underwent electrophysiological recordings from ventral striatum during the training session. To identify specific cell types, mice were previously injected with a Cre-dependent virus that drove expression of the excitatory opsin channelrhodopsin-2 (ChR2)^34^. Recordings were carried out using a silicon microprobe attached to an optical fiber for light delivery (**Fig. 2a**)^35^. D1 and D2 MSNs were identified using an optogenetic tagging (i.e., opto-tagging) protocol applied immediately after the training session^36^. Subsequent histological analysis confirmed the co-localization of tyrosine hydroxylase (TH) expressing dopamine terminals, ChR2, and the silicon microprobe in the nucleus accumbens region of ventral striatum (**Fig. 2b**). In line with previous publications^22–24,37,38^, one of the opto-tagging criteria was short latency neuronal activation to blue light presentation. Using the well-established maximum latency criterion of 6 ms, 95 D1 and 115 D2 MSNs were recorded from a total of 9 D1-Cre and 8 A2a-Cre mice. The primary analysis was carried out on these cell populations. Applying a shorter maximum latency resulted in fewer opto-tagged cells (**Fig. 2c**). Despite this constraint, the main results of this study were confirmed using an even more stringent maximum latency criterion of 3 ms (**Extended Data Fig. 2**). **Fig. 2d** shows an example of an opto-tagged D1 MSN with rapid activation following the laser and a spike waveform that was unaltered by laser presentation. The same cell exhibited increased firing to both cues, but the response was stronger on CS+ relative to CS- trials (**Fig. 2e**). Consequently, this cell was classified as being more selective for CS+ (**Fig. 2f**, selectivity index range is ±1, with positive/negative values indicating greater CS+/CS- activation). The neural modulation and selectivity to cues varied qualitatively across the four 25-trial blocks, suggesting altered stimulus representations as a function of learning. A subset of D2 MSNs also exhibited firing properties that appeared to change over the course of the training session (example shown in **Fig. 2g-i**). While these example cells demonstrated that the opto-tagging technique could be successfully applied during learning, they did not necessarily reflect the average properties of each cell type nor how each population’s activity related to behavior.

**Fig. 2.**
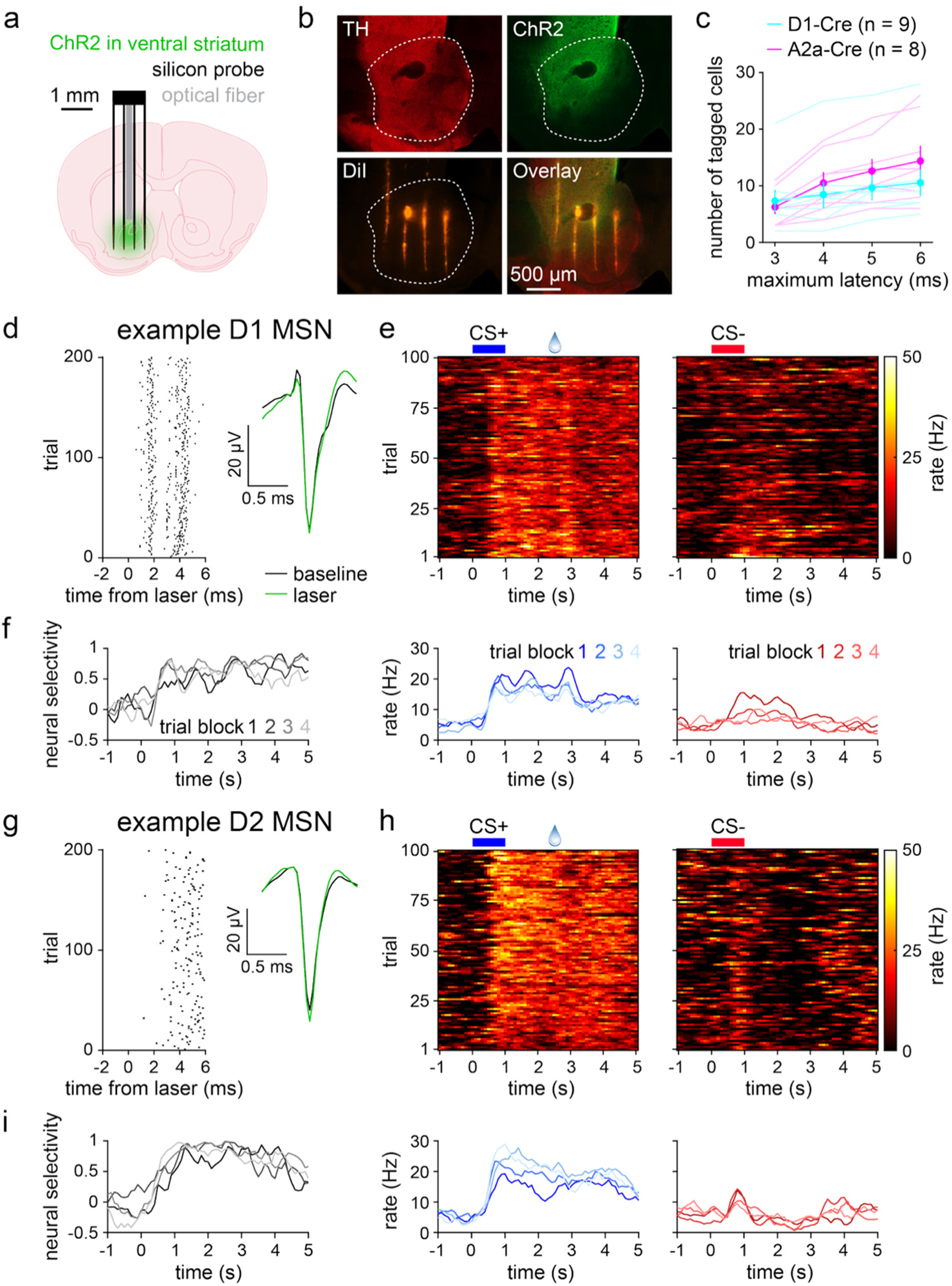
Electrophysiological recordings of ventral striatal D1 and D2 MSNs. **a.** Experimental approach for electrophysiological recording of opto-tagged D1 and D2 MSNs. **b.** Fluorescence image of a brain section from an A2a-Cre mouse depicting TH-expressing dopaminergic neuron terminals, ChR2 in D2 MSNs, DiI stain from the silicon microprobe, and image overlay. Dashed line indicates the region of ventral striatum that was targeted for recordings. **c.** Number of tagged MSNs per animal versus maximum latency for identifying tagged cells (*n* = 9 D1-Cre and 8 A2a-Cre mice; two-way RM ANOVA; latency effect: *F*_3,45_ = 33, *P* < 0.0001; cell type effect: *F*_1,15_ = 0.5, *P* = 0.50). Dark lines and error bars depict mean ± SEM; light lines depict individual animal data. **d.** Laser-evoked spike raster (left) and spike waveforms (right) for a D1 MSN. **e.** D1 MSN activity during the 100 CS+ trials (top left), 100 CS- trials (top right), and averaged across the 25 trials of each block (bottom). **f.** D1 MSN neural selectivity averaged within trial blocks. d-f depict data from the same D1 MSN. **g-i.** Same as d-f, but for a D2 MSN. See also Extended Data Fig. 2.

### D1 MSN selectivity increases with learning and correlates with behavior

We next determined how D1 and D2 MSN stimulus discrimination varied across learning and individual animals. As with the behavioral data, the training session was divided into four blocks of 25 trials of each stimulus type. Neural selectivity was evaluated in the interval between cue onset and reward on CS+ trials and an equal duration interval on CS- trials. Both D1 and D2 MSNs exhibited learning-dependent changes in cue discrimination, but to markedly different degrees. For D1 MSNs, the mean percentage of cue-selective cells and the neural selectivity index increased significantly across the four blocks (**Fig. 3a-c**). The positive neural selectivity indicated a higher response on CS+ relative to CS- trials. In addition, both measures of D1 MSN cue selectivity were significantly correlated with lick selectivity, suggesting a strong coupling between D1 MSN and behavioral discrimination (**Fig. 3d,e**). By contrast, D2 MSNs showed a statistically significant change in the percentage of cue-selective cells, but no significant change in mean neural selectivity (**Fig. 3f-h**). Furthermore, for D2 MSNs, neither of these parameters were correlated with lick selectivity, implying a relatively weak relationship between D2 MSN activity and behavior (**Fig. 3i,j**). Thus, the stimulus discrimination properties of D1 and D2 MSNs were differentially regulated during learning. Namely, learning was primarily associated with increased D1 MSN selectivity, and these neurophysiological properties were correlated with individual animal performance on the stimulus discrimination task.

**Fig. 3.**
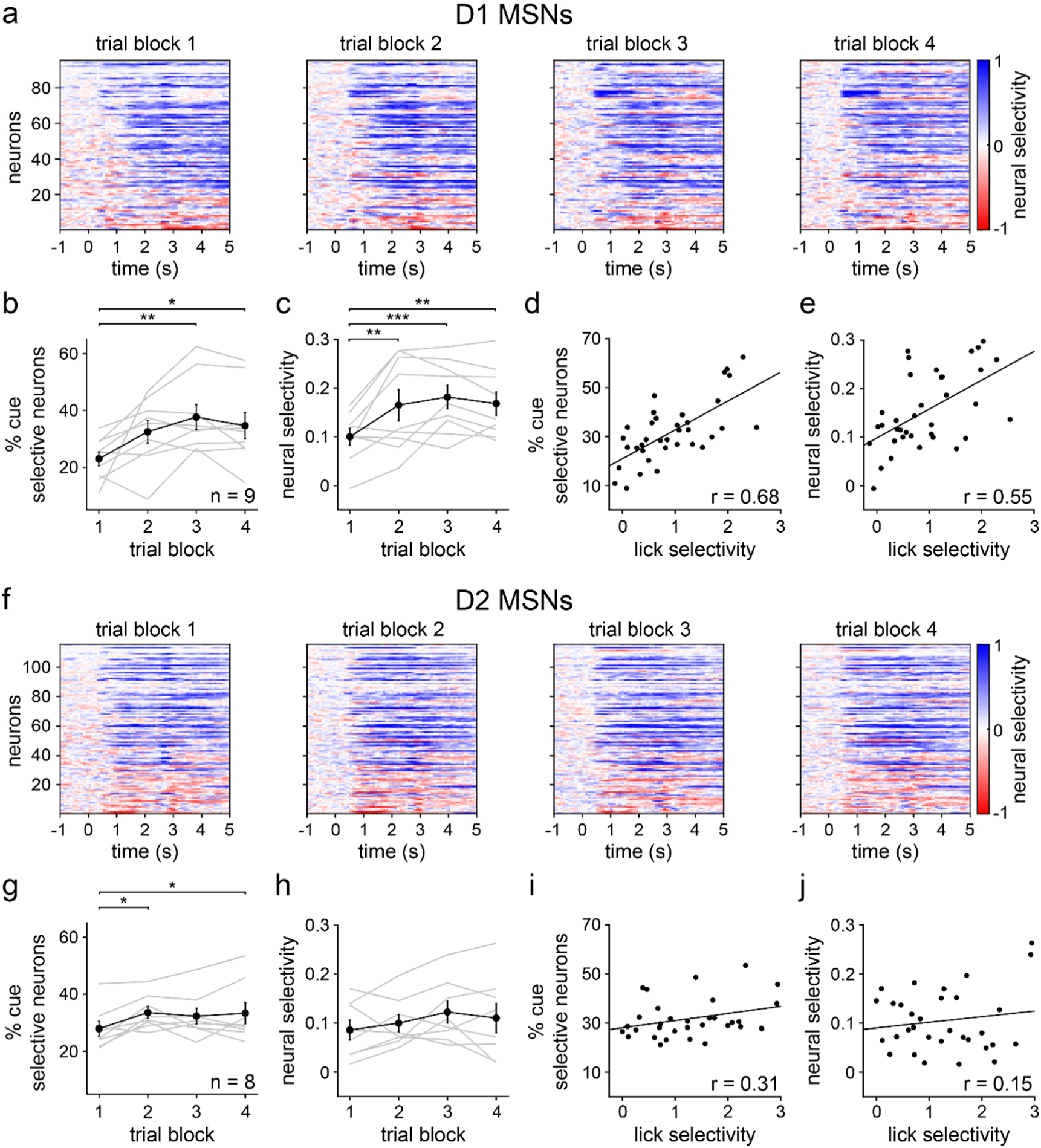
D1 MSN selectivity increases with learning and correlates with lick selectivity. **a.** Trial-averaged neural selectivity of D1 MSNs for each of the four 25-trial blocks. Positive/negative values indicate a larger/smaller response to CS+ relative to CS-. **b.** Percentage of D1 MSNs selectively responsive to CS+ or CS- versus trial block (*n* = 9 D1-Cre mice; one-way RM ANOVA; block effect: *F*_3,24_ = 5, *P* = 0.0069; multiple comparisons for blocks 1 versus 2 (*P* = 0.062), 3 (*P* = 0.0030), and 4 (*P* = 0.019)). **c.** D1 MSN selectivity versus trial block (one-way RM ANOVA; block effect: *F*_3,24_ = 8, *P* = 0.0006; multiple comparisons for blocks 1 versus 2 (*P* = 0.0040), 3 (*P* = 0.0004), and 4 (*P* = 0.0026)). **d.** Percentage of cue selective D1 MSNs versus lick selectivity (*n* = 36 (9 mice and 4 trial blocks); Pearson correlation coefficient; *P* < 0.0001). **e.** D1 MSN selectivity versus lick selectivity (Pearson correlation coefficient; *P* = 0.0005). **f.** Same as a, but for D2 MSNs. **g.** Same as b, but for D2 MSNs (*n* = 8 A2a-Cre mice; one-way RM ANOVA; block effect: *F*_3,21_ = 4, *P* = 0.017; multiple comparisons for blocks 1 versus 2 (*P* = 0.014), 3 (*P* = 0.062), and 4 (*P* = 0.018)). **h.** Same as c, but for D2 MSNs (one-way RM ANOVA; block effect: *F*_3,21_ = 1.4, *P* = 0.27). Dark colors in b-c and g-h depict mean ± SEM; light colors depict individual animal data. **i.** Same as d, but for D2 MSNs (*n* = 32 (8 mice and 4 trial blocks); Pearson correlation coefficient; *P* = 0.088). **j.** Same as e, but for D2 MSNs (Pearson correlation coefficient; *P* = 0.41). * *P* < 0.05, ** *P* < 0.01, *** *P* < 0.001.

### Average CS+ evoked D1 and D2 MSN activity does not change with learning

The increase in D1 MSN selectivity may result either from learning-dependent increases in CS+ responses, decreases in CS- responses, or both. We sought to clarify these possible mechanisms, starting with CS+ evoked activity. A sizable subset of D1 and D2 MSNs responded to reward-predictive cues within hundreds of milliseconds of odor presentation (**Fig. 4a,b**). However, at the population level, the mean modulation indices exhibited inconsistent, statistically insignificant changes across learning (**Fig. 4c,d**, modulation index range is ±1, with positive/negative values indicating cue-evoked excitation/inhibition relative to baseline). Likewise, the average percentage of CS+ modulated cells displayed insignificant changes (**Fig. 4e,f**). To rule out the possibility of positive and negative CS+ responses canceling out, we subdivided cells into three groups according to their response properties (excited, inhibited, and non-modulated, **Extended Data Fig. 3a,b**). Overall, D1 and D2 MSNs exhibited mixed responses to the CS+, though most cells were classified as excited (65% for D1 and 54% for D2 MSNs). Several mice contained zero opto-tagged cells that were inhibited or not modulated by cues. Thus, **Extended Data Fig. 3** analyses were conducted by pooling cells across animals, rather than on a per animal basis, which was used throughout the rest of this study. In general, all three response groups showed weak changes in CS+ evoked firing as a function of block number, except for a statistically significant change in the D1 MSN inhibited group (**Extended Data Fig. 3c-h**). However, since the inhibitory response showed a small increase with learning (i.e., the mean modulation index became more negative), this cannot explain the overall increase in D1 MSN cue selectivity.

**Fig. 4.**
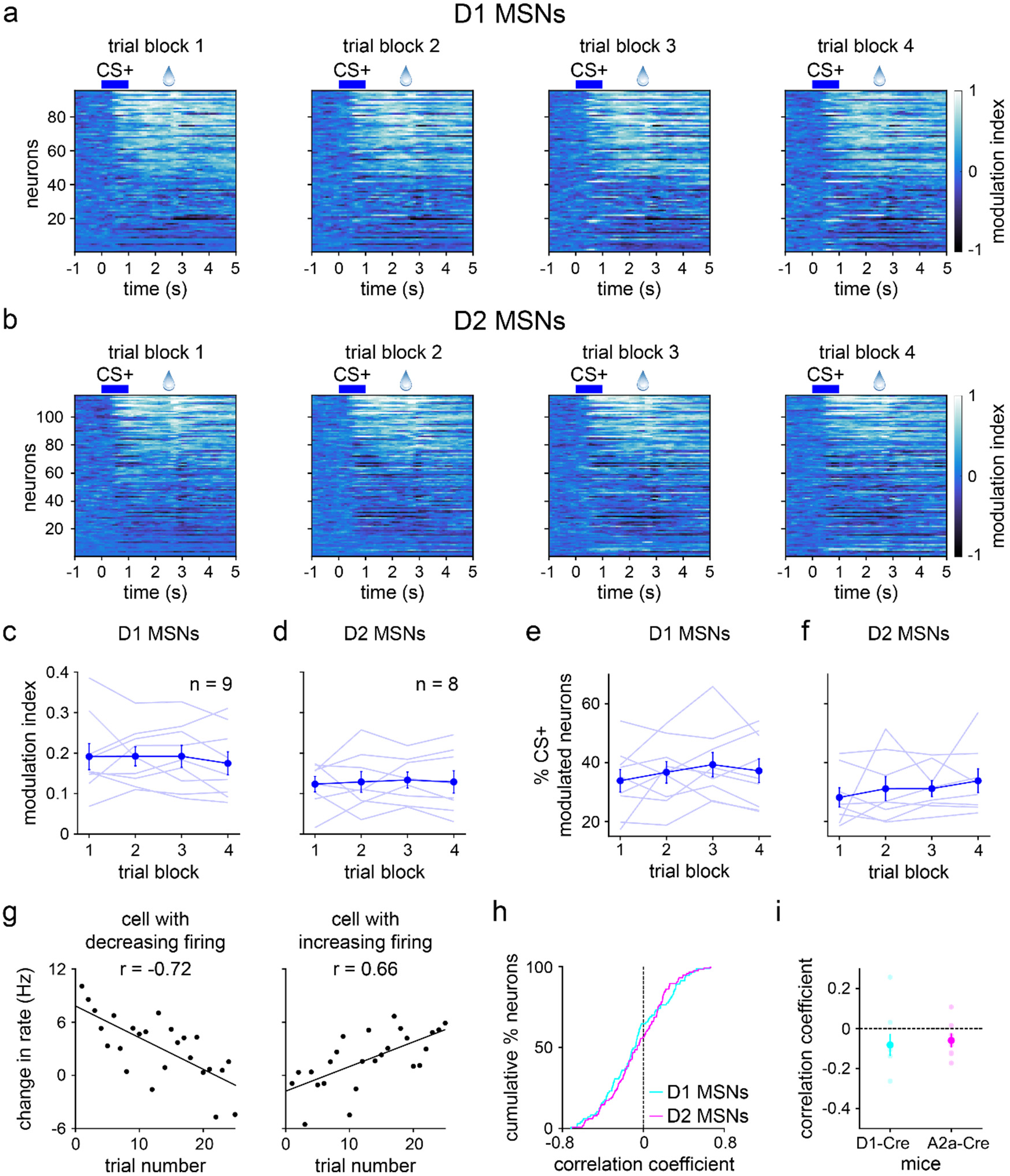
Average CS+ evoked D1 and D2 MSN activity does not change with learning. **a.** Trial-averaged modulation index of D1 MSNs for each of the four 25-trial blocks. Positive/negative values indicate increased/decreased response to CS+ relative to baseline. **b.** Same as a, but for D2 MSNs. **c.** D1 MSN modulation index versus trial block (*n* = 9 D1-Cre mice; one-way RM ANOVA; block effect: *F*_3,24_ = 0.3, *P* = 0.80). **d.** Same as c, but for D2 MSNs (*n* = 8 A2a-Cre mice; one-way RM ANOVA; block effect: *F*_3,21_ = 0.09, *P* = 0.96). **e.** Percentage of CS+ modulated D1 MSNs versus trial block (*n* = 9 D1-Cre mice; one-way RM ANOVA; block effect: *F*_3,24_ = 0.9, *P* = 0.46). **f.** Same as e, but for D2 MSNs (*n* = 8 A2a-Cre mice; one-way RM ANOVA; block effect: *F*_3,21_ = 0.9, *P* = 0.47). **g.** Baseline-subtracted firing rate versus trial number (first 25 trials) for two D1 MSNs. With learning, one cell decreases activity in response to CS+ (*n* = 25 trials; Pearson correlation coefficient; *P* < 0.0001), and another cell increases activity to CS+ (Pearson correlation coefficient; *P* = 0.0003). **h.** Cumulative percentage of D1 and D2 MSNs versus correlation coefficient between firing rate and trial number (first 25 trials) (*n* = 95 D1 and 115 D2 MSNs; two-sample Kolmogorov-Smirnov test; *P* = 0.68). **i.** Correlation coefficient between firing rate and trial number (first 25 trials), plotted for each MSN type (*n* = 9 D1-Cre and 8 A2a-Cre mice; two-sample t-test; *P* = 0.73). Dark colors in c-f and i depict mean ± SEM; light colors depict individual animal data. See also Extended Data Fig. 3-5.

These results indicate a surprisingly static neural representation of reward-predictive cues across the four training blocks. However, this does not rule out the presence of robust changes within individual blocks. Indeed, as already seen with behavior (**Extended Data Fig. 1d,e**), D1 MSN firing properties changed appreciably within the first 25-trial block of training, though D2 MSN activity remained unaltered (**Extended Data Fig. 4**). To investigate whether CS+ responses were rapidly altered in the early training phase, we focused on the initial 25 trials and assessed cue-evoked firing properties on a trial-by-trial basis. Both MSN populations showed heterogeneous firing rate changes, with different cells increasing, decreasing, or maintaining activity as a function of trial number (**Fig. 4g** and **Extended Data Fig. 5**). These trends were quantified by calculating the Pearson correlation between firing rate and trial number. On average, the correlation coefficients remained near zero and did not display significant cell type differences (**Fig. 4h,i**). Thus, echoing our qualitative observations, a subset of both D1 and D2 MSNs did show learning-dependent changes in CS+ evoked activity. Crucially, however, these effects were mixed and lacked a consistent directional trend at the average population level. Furthermore, D1 and D2 MSNs did not markedly differ in how their CS+ responses varied across learning. Therefore, it appears unlikely that the observed increase in D1 MSN cue selectivity across learning is driven by altered CS+ evoked activity.

It has been reported that learning alters reward-evoked striatal neuronal activity^30,39^, although other work suggests a learning-independent, innate response to appetitive stimuli^11^. Our analysis revealed no significant changes in D1 and D2 MSN modulation in the period immediately after reward delivery (**Extended Data Fig. 6**). Together, these findings suggest, on average, a relatively fixed representation of reward-predictive cues as well as the rewards themselves during stimulus discrimination learning.

### Learning attenuates CS- evoked D1 MSN responses

Since CS+ evoked activity changed little on average across learning, we hypothesized that the improvement in neural stimulus discrimination emerges from changes in CS- evoked responses. Consistent with this idea, a reduction in CS- evoked activity across trial blocks was observed for D1 MSNs and was markedly less pronounced for D2 MSNs (**Fig. 5a,b**). Both the mean modulation index and percentage of CS- modulated neurons significantly decreased across learning for D1, but not D2, MSNs (**Fig. 5c-f**). Subdividing cells by their response properties, the main learning-dependent change was again an attenuation effect, found in D1 MSNs excited by CS- (**Extended Data Fig. 3i-p**). We also observed a significant increase in the inhibitory response of D2 MSNs (i.e., increasingly negative mean modulation index), but this effect was comparatively small. To confirm that the attenuation of D1 MSN responses was also robustly observed in the early phase of training, we assessed the trial-by-trial firing rate during the initial 25 trials. Both MSN populations contained cells with either increasing, decreasing, or fixed CS- evoked responses as a function of trial number (**Fig. 5g** and **Extended Data Fig. 5**). However, on average, correlations between D1 MSN firing rate and trial number were biased toward significantly more negative values than for D2 MSNs (**Fig. 5h,i**). Collectively, these findings indicate that learning a stimulus discrimination task is accompanied by elevated D1 MSN cue selectivity, and these altered neural activity patterns are primarily driven by reduced CS- evoked excitation.

**Fig. 5.**
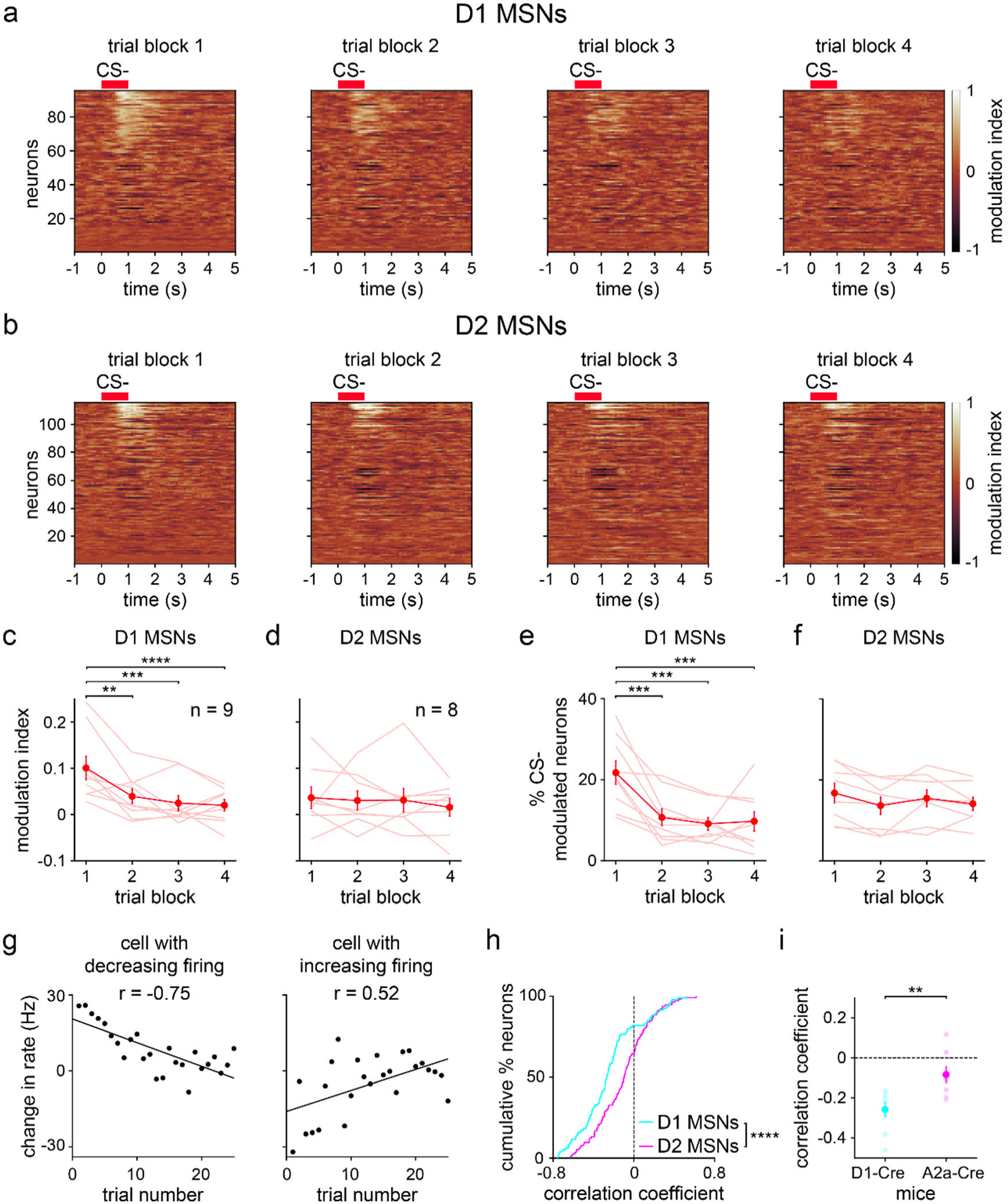
Average CS- evoked D1 MSN activity decreases during learning. **a.** Trial-averaged modulation index of D1 MSNs for each of the four 25-trial blocks. Positive/negative values indicate increased/decreased response to CS- relative to baseline. **b.** Same as a, but for D2 MSNs. **c.** D1 MSN modulation index versus trial block (*n* = 9 D1-Cre mice; one-way RM ANOVA; block effect: *F*_3,24_ = 11, *P* < 0.0001; multiple comparisons for blocks 1 versus 2 (*P* = 0.0019), 3 (*P* = 0.0002), and 4 (*P* < 0.0001)). **d.** Same as c, but for D2 MSNs (*n* = 8 A2a-Cre mice; one-way RM ANOVA; block effect: *F*_3,21_ = 0.3, *P* = 0.81). **e.** Percentage of CS- modulated D1 MSNs versus trial block (*n* = 9 D1-Cre mice; one-way RM ANOVA; block effect: *F*_3,24_ = 10, *P* = 0.0001; multiple comparisons for blocks 1 versus 2 (*P* = 0.0008), 3 (*P* = 0.0002), and 4 (*P* = 0.0003)). **f.** Same as e, but for D2 MSNs (*n* = 8 A2a-Cre mice; one-way RM ANOVA; block effect: *F*_3,21_ = 2, *P* = 0.15). **g.** Baseline-subtracted firing rate versus trial number (first 25 trials) for two D1 MSNs. With learning, one cell decreases activity in response to CS- (*n* = 25 trials; Pearson correlation coefficient; *P* < 0.0001), and another cell increases activity to CS- (Pearson correlation coefficient; *P* = 0.0078). **h.** Cumulative percentage of D1 and D2 MSNs versus correlation coefficient between firing rate and trial number (first 25 trials) (*n* = 95 D1 and 115 D2 MSNs; two-sample Kolmogorov-Smirnov test; *P* < 0.0001). **i.** Correlation coefficient between firing rate and trial number (first 25 trials), plotted for each MSN type (*n* = 9 D1-Cre and 8 A2a-Cre mice; two-sample t-test; *P* = 0.0043). Dark colors in c-f and i depict mean ± SEM; light colors depict individual animal data. ** *P* < 0.01, *** *P* < 0.001, **** *P* < 0.0001. See also Extended Data Fig. 3-5.

### D1 MSN activation on CS- trials impairs stimulus discrimination

Finally, we examined the potential behavioral relevance of the changes in neural activity observed during learning. Since the most salient change was an attenuation in CS- evoked D1 MSN activity, we tested the hypothesis that activating D1 MSNs selectively on CS- trials would impair learned stimulus discrimination. A separate group of D1-Cre mice was injected with ChR2 and implanted with permanent optical fibers unilaterally in ventral striatum (**Fig. 6a,b**). Mice were pre-trained to perform the same stimulus discrimination task described thus far in the absence of optogenetic stimulation. Animals then underwent a behavioral testing session in which CS- trials were consistently paired with optogenetic stimulation for a duration of 2.5 s (**Fig. 6c**). CS+ trials were not paired with laser and continued to be randomly interleaved with CS- trials. Behavior on the day before stimulation exhibited robust anticipatory licking to the CS+ and sparse licking to the CS- (**Fig. 6d**). D1 MSN activation markedly increased licking to the CS- (**Fig. 6e**). As expected based on the electrophysiological results, this manipulation significantly reduced the lick selectivity of ChR2 expressing animals, reflecting an increase in CS- evoked lick probability (**Fig. 6f,g**). Notably, however, a significant decrease in CS+ evoked lick probability was concomitantly observed, even though optogenetic stimulation was never temporally associated with this cue (**Fig. 6h**). Therefore, stimulus discrimination was impaired by two mutually adverse effects, both higher CS- licking and lower CS+ licking. Together with the behaviorally correlated neural activity (**Fig. 3d,e**), these findings bolster support for D1 MSNs causally contributing to discrimination between reward-paired and unpaired stimuli.

**Fig. 6.**
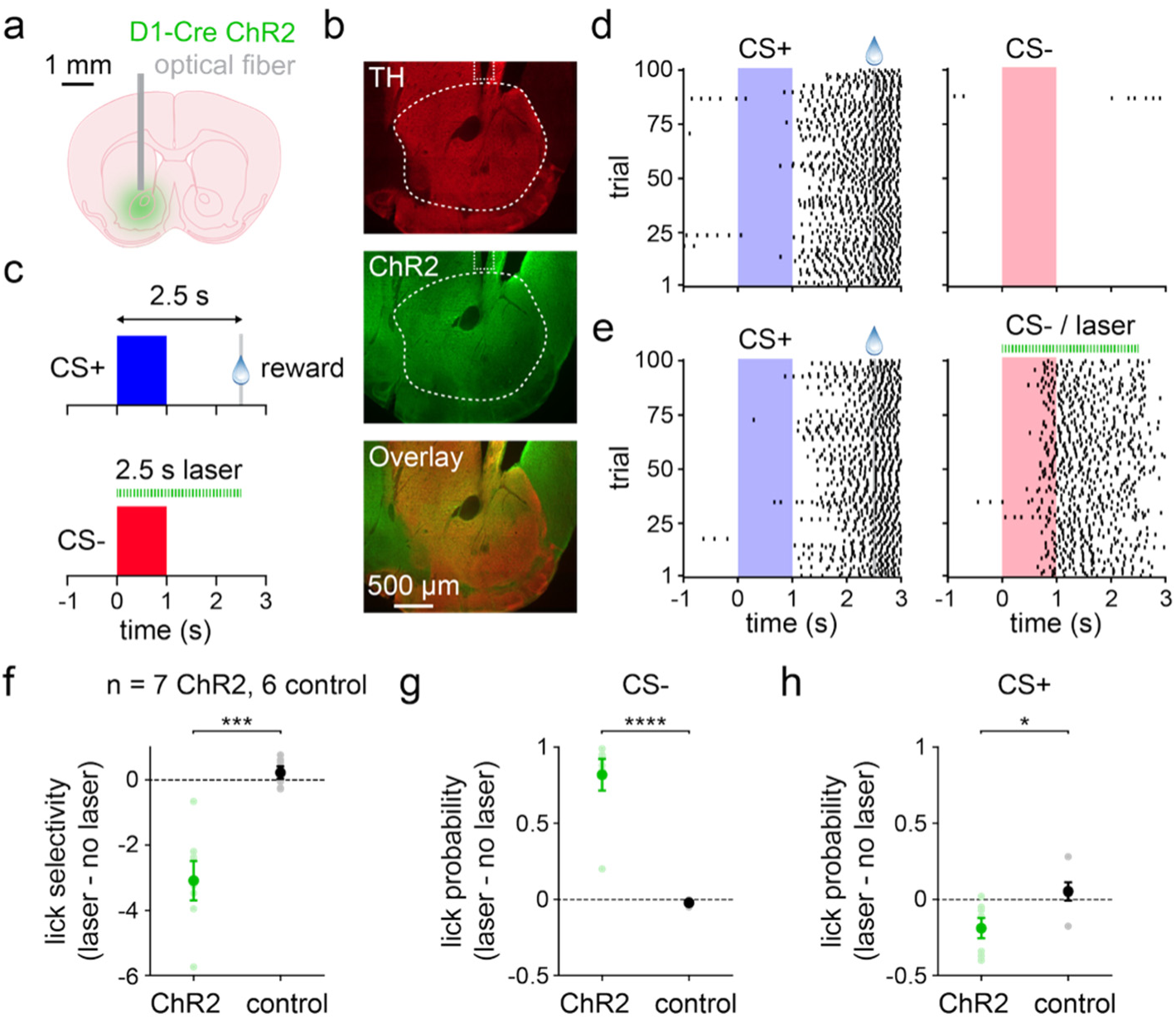
D1 MSN activation during CS- trials impairs stimulus discrimination. **a.** Experimental approach for optogenetic activation of D1 MSNs. **b.** Histology from a D1-Cre mouse depicting TH-expressing dopaminergic neuron terminals, ChR2 in D1 MSNs, and image overlay. Dashed lines indicate approximate location of the optical fiber tip and the targeted region of ventral striatum. **c.** Schematic of the discrimination task. Mice pre-trained on the CS+ / CS- task with no laser stimulation (not depicted) then trained on a similar task with laser-paired CS-. **d.** Lick rasters from one mouse on CS+ (left) and CS- (right) trials. **e.** Lick rasters from the same mouse on CS+ (left) and laser-paired CS- (right) trials. **f.** Differences in lick selectivity between laser and no laser training days for ChR2 and control mice (*n* = 7 ChR2 and 6 control mice; two-sample t-test; *P* = 0.0005). **g.** Same as f, but for CS- lick probability (two-sample t-test; *P* < 0.0001). **h.** Same as f, but for CS+ lick probability (two-sample t-test; *P* = 0.022). Dark colors in f-h depict mean ± SEM; light colors depict individual animal data. * *P* < 0.05, *** *P* < 0.001, **** *P* < 0.0001.

## DISCUSSION

Based on a lack of direct prior evidence, we presented several potential models for how learning could theoretically impact the cue discrimination properties of ventral striatal D1 and D2 MSNs (**Fig. 1a**). Based on electrophysiological recordings from optogenetically identified neurons, our results are most consistent with the first model: the main effect of stimulus discrimination learning is an enhancement in D1 MSN selectivity. The selectivity of D1 MSNs was significantly correlated with behavioral performance on the stimulus discrimination task. In contrast, D2 MSNs displayed less dynamic, behaviorally uncorrelated activity over the course of learning. These findings indicate markedly different effects of stimulus discrimination learning on D1 and D2 MSN firing properties. This does not imply that one cell type is more functionally important than the other; D2 MSNs are likely to play a role in discriminative behaviors^40^, but their contributions may be less dependent on learning.

A limited amount of work has previously examined the impact of reward-guided learning on D1 and D2 MSN activity specifically in ventral striatum^29,30^. Interestingly, our results appear at odds with the main conclusions of the study by Zachry and colleagues^30^, whose observations were in a diametrically opposite direction. Using operant conditioning tasks, the authors showed that D1 MSN activity evoked by reward- or shock-paired cues does not appreciably change with learning, while D2 MSN activity increases significantly. Because Zachry et al. did not use a stimulus discrimination task with reward-paired and unpaired cues, a direct comparison with our findings is not possible. Nevertheless, this potential discrepancy may reflect methodological differences in the behavioral tasks (classical versus operant conditioning), striatal subregions (ventral striatum versus nucleus accumbens core), or recording techniques (electrophysiology versus calcium imaging)^41^, all of which warrant further investigation.

Surprisingly, in our study, the learning-mediated enhancement in D1 MSN selectivity is not strongly attributed to an increased excitatory response to rewarded cues (CS+). Instead, it is primarily driven by a reduced excitatory response to unrewarded cues (CS-). The most straightforward interpretation of this effect is that learning attenuates D1 MSN activity related to unrewarded stimuli in order to curtail unproductive behavioral responding (i.e., to reduce anticipatory licking to cues that are never followed by rewards). In agreement with this view, we observed that artificially raising D1 MSN activity during unrewarded cues is sufficient to significantly disrupt the normal pattern of behavioral stimulus discrimination. Collectively, these results provide further support for a role for ventral striatum in behavioral inhibition, an effect proposed to suppress impulsive or unproductive reward seeking^12–14^. However, previous work, including that of Ambroggi and colleagues, generally favors a model in which ventral striatum actively suppresses behavioral responding to unrewarded stimuli^13^. If this were the case, we would expect learning to be accompanied by an increased excitatory response to unrewarded cues. However, the opposite effect was observed: CS- evoked activity was attenuated in D1 MSNs. Thus, the main mechanism for behavioral inhibition in our task appears to be a learning-dependent reduction in lick-promoting signals from D1 MSNs, a phenomenon more analogous to suppressing movement by releasing the gas pedal rather than stepping on the brakes.

A further intriguing implication of these results is that the initially high and indiscriminate level of D1 MSN cue-evoked activity may promote exploration of multiple different stimuli. Indeed, an early exploratory phase of learning may facilitate the acquisition of appropriate actions contingent on different sensory cues^42^. From this perspective, our findings suggest that ventral striatal D1 MSNs serve a disproportionately important role in the early stage of learning by enabling exploration of the stimulus-motor response space^43^. We found that, as the reward contingencies in the task are better established later in learning, CS- evoked D1 MSN activity is attenuated, thereby mitigating further unnecessary exploratory behavior.

While the observed changes in CS- evoked neural activity can explain the reduced licking to unrewarded cues, the relationship between CS+ evoked activity and behavior during learning is less apparent. Notably, while animals greatly increased CS+ evoked licking across the training session, on average, neither D1 nor D2 MSNs displayed consistent changes in CS+ evoked activity. Thus, a major question is whether the neural activity patterns that we observed can potentially explain the increased licking response to reward-paired cues. Individual cells displayed a wide range of alterations in cue-evoked firing rate over the course of the training session. These mixed positive and negative changes largely canceled out when averaging across the population. Thus, at the population level both cell types displayed relatively fixed CS+ responses. On one hand, it is conceivable that downstream brain areas do not read out a simple linear average of population activity, but a nonlinear weighted sum of individual neural responses^44^. Such a mechanism could theoretically enable animals to respond preferentially to D1 MSNs with increased CS+ evoked firing during learning. On the other hand, it would be puzzling if learning were characterized by complex dynamical representations of reward-paired cues, but relatively straightforward alterations in unrewarded cue representations by the same cells. Thus, an alternative interpretation is that ventral striatum is not the primary region responsible for the learning-dependent increase in licking to rewarded cues. Other circuits known to contribute to motor learning, including dorsal striatum, are well suited to serve this function^39,45–48^. However, it is not yet clear whether the potentially limited role of ventral striatum in learning to lick to reward-predictive cues extends to other behavioral tasks and reinforcers^25,30^.

An additional possibility is suggested from the behavioral changes induced by optogenetically activating D1 MSNs. The main effect of D1 MSN activation during the unrewarded cue was an increase in licking on CS- trials. Interestingly, however, the same perturbations also caused a significant reduction in anticipatory licking evoked by the rewarded cue, even though neural activity was never manipulated during those trials. This indicates that CS- evoked D1 MSN activity is capable of suppressing behavior on subsequent trials containing reward-predictive cues. This suggests that the learning-dependent attenuation of CS- evoked D1 MSN responses may serve dual functions, reducing licking on unrewarded trials and disinhibiting licking on future rewarded trials. Taken together, these findings provide significant insight into the differential changes displayed by ventral striatal D1 and D2 MSNs as mice learn a stimulus discrimination task.

## EXTENDED DATA

**Extended Data Fig. 1.**
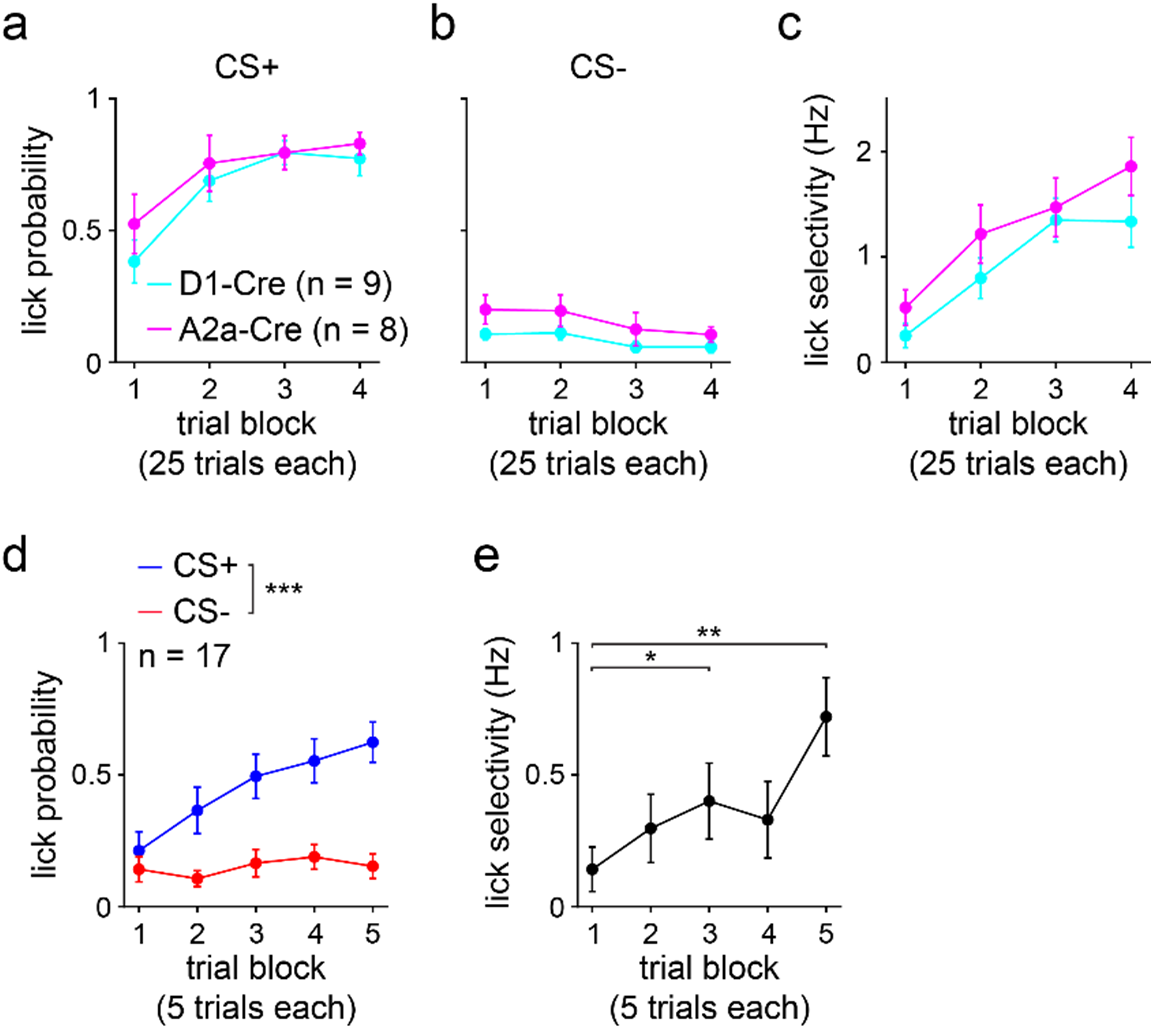
Behavior of D1-Cre and A2a-Cre mice and initial responses to cues. **a.** CS+ lick probability plotted separately for D1-Cre and A2a-Cre mice (*n* = 9 D1-Cre and 8 A2a-Cre mice; two-way RM ANOVA; block effect: *F*_3,45_ = 15, *P* < 0.0001; group effect: *F*_1,15_ = 0.7, *P* = 0.43). **b.** Same as a, but for CS- (two-way RM ANOVA; block effect: *F*_3,45_ = 4, *P* = 0.022; cell type effect: *F*_1,15_ = 3, *P* = 0.11). **c.** Same as a, but for lick selectivity (two-way RM ANOVA; block effect: *F*_3,45_ = 21, *P* < 0.0001; cell type effect: *F*_1,15_ = 1.8, *P* = 0.19). **d.** CS+ and CS- lick probabilities across the initial five 5-trial blocks (*n* = 17 mice; two-way RM ANOVA; block effect: *F*_4,128_ = 9, *P* < 0.0001; cue effect: *F*_1,32_ = 16, *P* = 0.0004; multiple comparisons for CS+ versus CS- on blocks 1 (*P* = 0.95), 2 (*P* = 0.030), 3 (*P* = 0.0025), 4 (*P* = 0.0006), and 5 (*P* < 0.0001)). **e.** Lick selectivity versus trial block (one-way RM ANOVA; block effect: *F*_3,42_ = 5, *P* = 0.0057; multiple comparisons for blocks 1 versus 2 (*P* = 0.37), 3 (*P* = 0.025), 4 (*P* = 0.51), and 5 (*P* = 0.0017)). a-e depict mean ± SEM. * *P* < 0.05, ** *P* < 0.01, *** *P* < 0.001.

**Extended Data Fig. 2.**
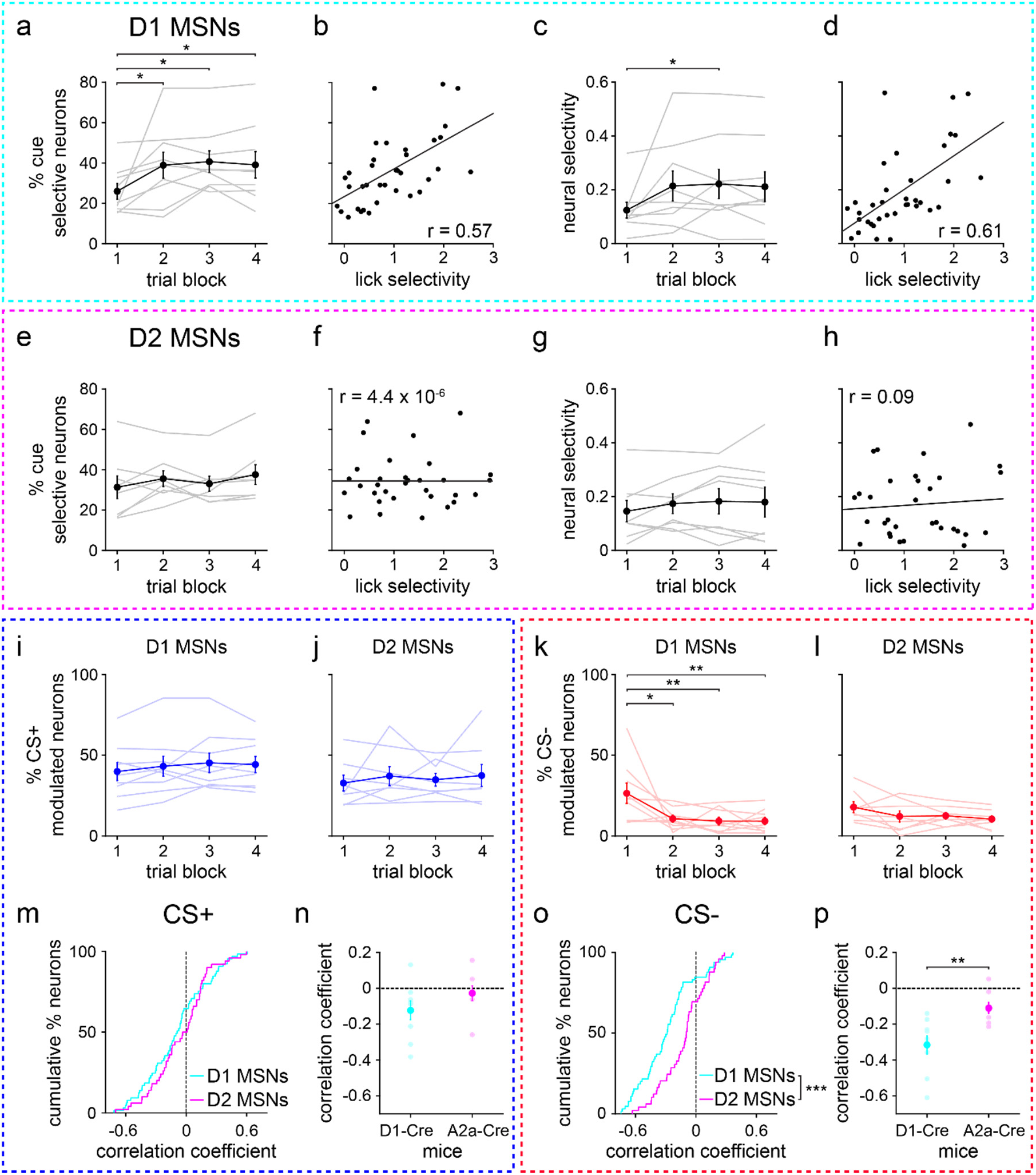
More stringent opto-tagging criteria replicate the main results. **a.** Percentage of D1 MSNs selectively responsive to CS+ or CS- versus trial block (*n* = 9 D1-Cre mice; one-way RM ANOVA; block effect: *F*_3,24_ = 4, *P* = 0.018; multiple comparisons for blocks 1 versus 2 (*P* = 0.033), 3 (*P* = 0.014), and 4 (*P* = 0.030)). **b.** Percentage of cue selective D1 MSNs versus lick selectivity (*n* = 36 (9 mice and 4 trial blocks); Pearson correlation coefficient; *P* = 0.0003). **c.** D1 MSN selectivity versus trial block (one-way RM ANOVA; block effect: *F*_3,24_ = 3, *P* = 0.041; multiple comparisons for blocks 1 versus 2 (*P* = 0.053), 3 (*P* = 0.034), and 4 (*P* = 0.062)). **d.** D1 MSN selectivity versus lick selectivity (Pearson correlation coefficient; *P* < 0.0001). **e.** Same as a, but for D2 MSNs (*n* = 8 A2a-Cre mice; one-way RM ANOVA; block effect: *F*_3,21_ = 1.7, *P* = 0.20). **f.** Same as b, but for D2 MSNs (*n* = 32 (8 mice and 4 trial blocks); Pearson correlation coefficient; *P* > 0.99). **g.** Same as c, but for D2 MSNs (one-way RM ANOVA; block effect: *F*_3,21_ = 1.1, *P* = 0.36). **h.** Same as d, but for D2 MSNs (Pearson correlation coefficient; *P* = 0.64). **i.** Percentage of CS+ modulated D1 MSNs versus trial block (one-way RM ANOVA; block effect: *F*_3,24_ = 1.1, *P* = 0.37). **j.** Percentage of CS+ modulated D2 MSNs versus trial block (one-way RM ANOVA; block effect: *F*_3,21_ = 0.4, *P* = 0.78). **k.** Percentage of CS- modulated D1 MSNs versus trial block (one-way RM ANOVA; block effect: *F*_3,24_ = 6, *P* = 0.0039; multiple comparisons for blocks 1 versus 2 (*P* = 0.010), 3 (*P* = 0.0051), and 4 (*P* = 0.0050)). **l.** Percentage of CS- modulated D2 MSNs versus trial block (one-way RM ANOVA; block effect: *F*_3,21_ = 1.7, *P* = 0.20). **m.** Cumulative percentage of D1 and D2 MSNs versus correlation coefficient between CS+ firing rate and trial number (first 25 trials) (*n* = 66 D1 and 50 D2 MSNs; two-sample Kolmogorov-Smirnov test; *P* = 0.47). **n.** Correlation coefficient between CS+ firing rate and trial number (first 25 trials), plotted for each MSN type (*n* = 9 D1-Cre and 8 A2a-Cre mice; two-sample t-test; *P* = 0.18). **o.** Same as m, but for CS- (two-sample Kolmogorov-Smirnov test; *P* = 0.0001). **p.** Same as n, but for CS-(two-sample t-test; *P* = 0.0048). All panels depict data using the 3 ms maximum latency criterion for opto-tagging. Dark colors in a, c, e, g, i-l, n, p depict mean ± SEM; light colors depict individual animal data. * *P* < 0.05, ** *P* < 0.01.

**Extended Data Fig. 3.**
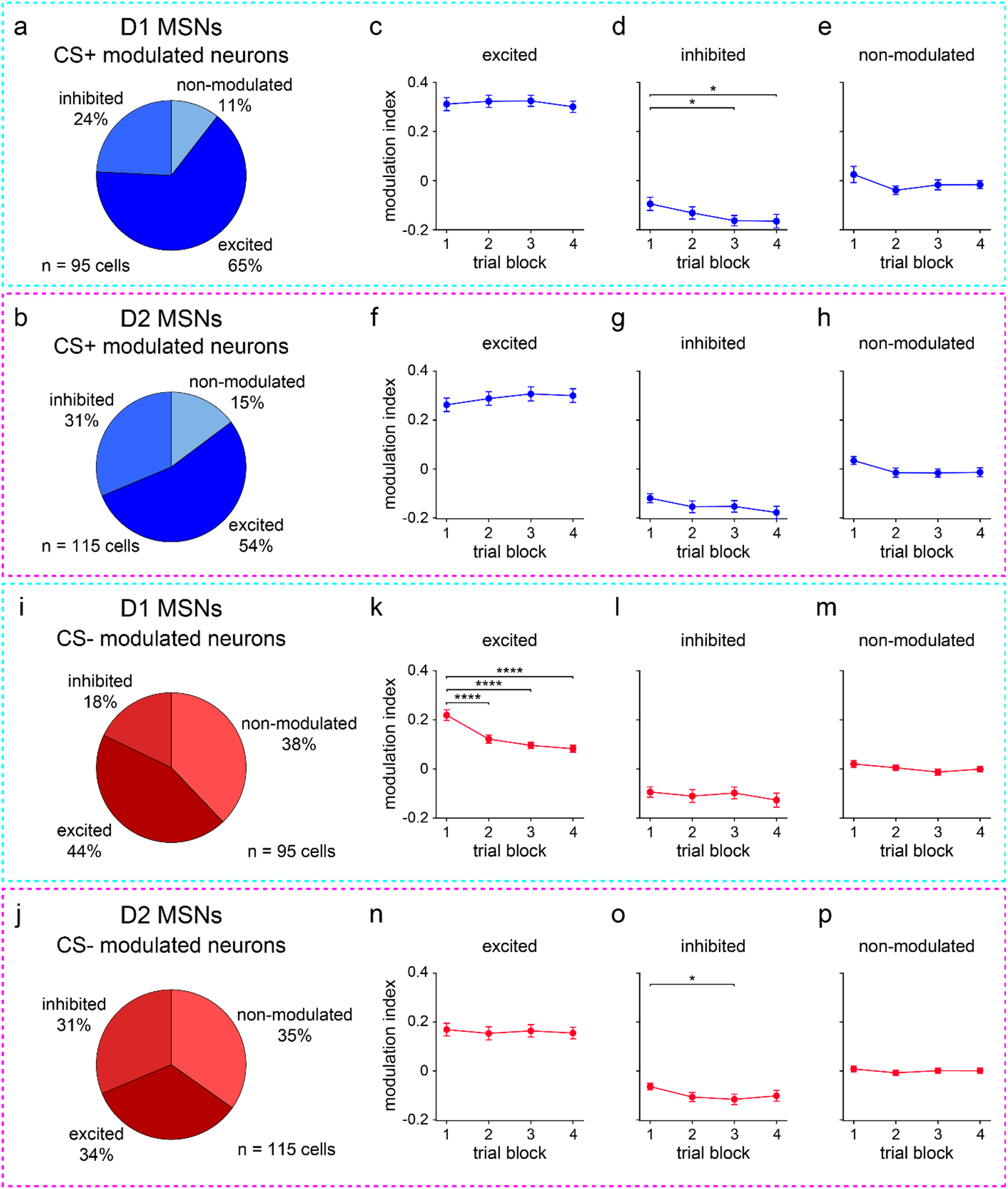
D1 and D2 MSNs exhibit mixed responses to predictive cues. **a.** Proportion of D1 MSNs that were excited, inhibited, or non-modulated by CS+. **b.** Same as a, but for D2 MSNs. **c.** Modulation index versus trial block for D1 MSNs excited by CS+ (*n* = 62 D1 MSNs; one-way RM ANOVA; block effect: *F*_3,183_ = 0.9, *P* = 0.44). **d.** Same as c, but for D1 MSNs inhibited by CS+ (*n* = 23 D1 MSNs; one-way RM ANOVA; block effect: *F*_3,66_ = 3, *P* = 0.031; multiple comparisons for blocks 1 versus 2 (*P* = 0.37), 3 (*P* = 0.032), and 4 (*P* = 0.026)). **e.** Same as c, but for D1 MSNs non-modulated by CS+ (*n* = 10 D1 MSNs; one-way RM ANOVA; block effect: *F*_3,27_ = 1.5, *P* = 0.23). **f.** Modulation index versus trial block for D2 MSNs excited by CS+ (*n* = 62 D2 MSNs; one-way RM ANOVA; block effect: *F*_3,183_ = 2, *P* = 0.11). **g.** Same as f, but for D2 MSNs inhibited by CS+ (*n* = 36 D2 MSNs; one-way RM ANOVA; block effect: *F*_3,105_ = 2, *P* = 0.086). **h.** Same as f, but for D2 MSNs non-modulated by CS+ (*n* = 17 D2 MSNs; one-way RM ANOVA; block effect: *F*_3,48_ = 2, *P* = 0.11). **i.** Proportion of D1 MSNs that were excited, inhibited, or non-modulated by CS-. **j.** Same as i, but for D2 MSNs. **k.** Modulation index versus trial block for D1 MSNs excited by CS- (*n* = 42 D1 MSNs; one-way RM ANOVA; block effect: *F*_3,123_ = 22, *P* < 0.0001; multiple comparisons for blocks 1 versus 2-4 (*P* < 0.0001)). **l.** Same as k, but for D1 MSNs inhibited by CS- (*n* = 17 D1 MSNs; one-way RM ANOVA; block effect: *F*_3,48_ = 0.7, *P* = 0.55). **m.** Same as k, but for D1 MSNs non-modulated by CS- (*n* = 36 D1 MSNs; one-way RM ANOVA; block effect: *F*_3,105_ = 1.3, *P* = 0.27). **n.** Modulation index versus trial block for D2 MSNs excited by CS- (*n* = 39 D2 MSNs; one-way RM ANOVA; block effect: *F*_3,114_ = 0.2, *P* = 0.87). **o.** Same as n, but for D2 MSNs inhibited by CS- (*n* = 36 D2 MSNs; one-way RM ANOVA; block effect: *F*_3,105_ = 3, *P* = 0.048; multiple comparisons for blocks 1 versus 2 (*P* = 0.082), 3 (*P* = 0.025), and 4 (*P* = 0.14)). **p.** Same as n, but for D2 MSNs non-modulated by CS- (*n* = 40 D2 MSNs; one-way RM ANOVA; block effect: *F*_3,117_ = 0.3, *P* = 0.80). c-h and k-p depict mean ± SEM. * *P* < 0.05, **** *P* < 0.0001.

**Extended Data Fig. 4.**
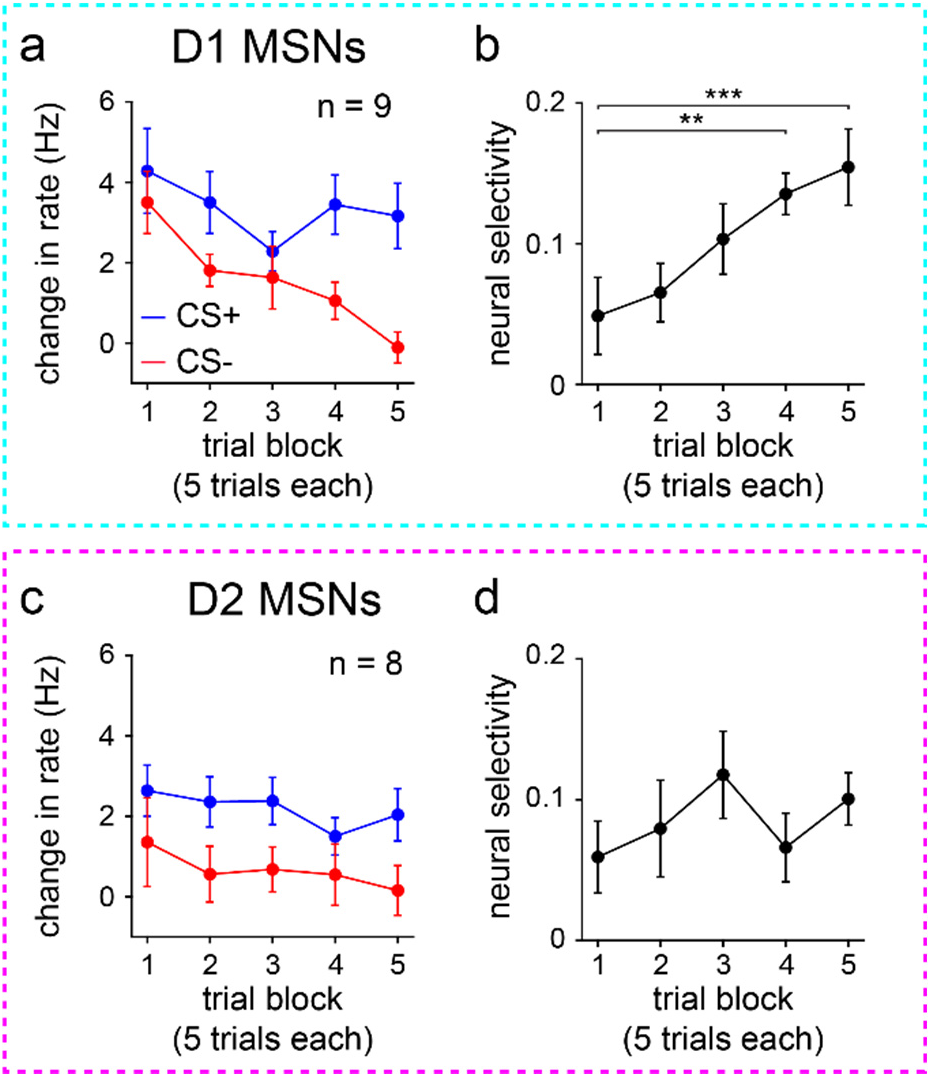
Changes in D1 MSN activity are evident early in learning. **a.** D1 MSN baseline-subtracted firing rate versus trial block (*n* = 9 D1-Cre mice; two-way RM ANOVA; block effect: *F*_4,64_ = 9, *P* < 0.0001; cue effect: *F*_1,16_ = 4, *P* = 0.051). **b.** D1 MSN selectivity versus trial block (one-way RM ANOVA; block effect: *F*_4,32_ = 6, *P* = 0.0009; multiple comparisons for blocks 1 versus 2 (*P* = 0.92), 3 (*P* = 0.13), 4 (*P* = 0.0068), and 5 (*P* = 0.0009)). **c.** Same as a, but for D2 MSNs (*n* = 8 A2a-Cre mice; two-way RM ANOVA; block effect: *F*_4,56_ = 1.3, *P* = 0.26; cue effect: *F*_1,14_ = 4, *P* = 0.067). **d.** Same as b, but for D2 MSNs (one-way RM ANOVA; block effect: *F*_4,28_ = 1.5, *P* = 0.23). a-d depict mean ± SEM. ** *P* < 0.01, *** *P* < 0.001.

**Extended Data Fig. 5.**
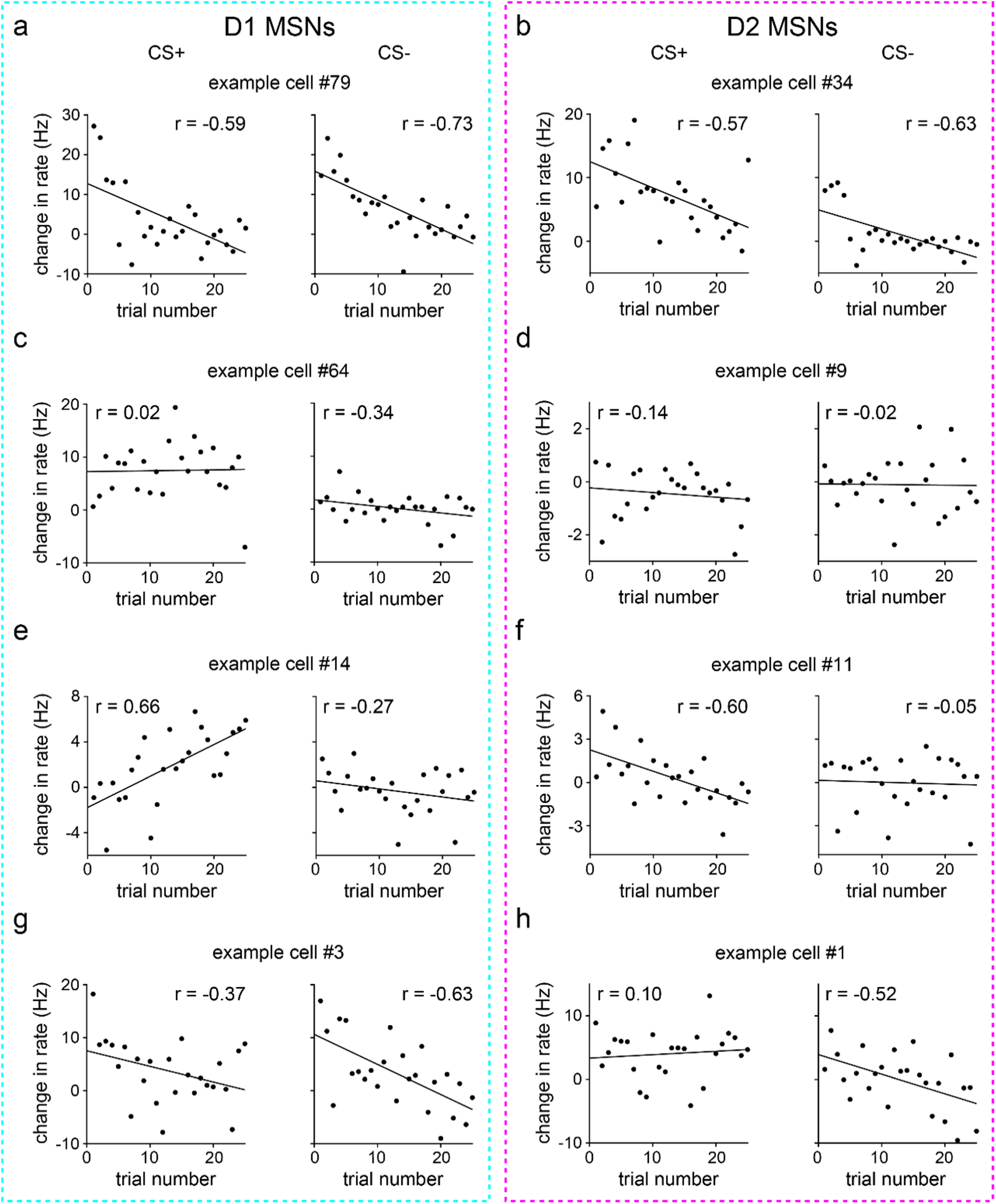
MSNs exhibit heterogenous changes in firing rate early in learning. **a.** D1 MSN with significantly decreasing firing rates to both CS+ and CS-. CS+ (*n* = 25 trials; Pearson correlation coefficient; *P* = 0.0018) and CS- (Pearson correlation coefficient; *P* < 0.0001). **b.** Same as a, but for D2 MSNs. CS+ (Pearson correlation coefficient; *P* = 0.0027) and CS-(Pearson correlation coefficient; *P* = 0.0007). **c.** D1 MSN with no significant change in firing rates to either CS+ or CS-. CS+ (Pearson correlation coefficient; *P* = 0.91) and CS- (Pearson correlation coefficient; *P* = 0.098). **d.** Same as c, but for D2 MSNs. CS+ (Pearson correlation coefficient; *P* = 0.50) and CS- (Pearson correlation coefficient; *P* = 0.93). **e.** D1 MSN with significant change in firing rates to CS+, but not CS-. CS+ (Pearson correlation coefficient; *P* = 0.0004) and CS-(Pearson correlation coefficient; *P* = 0.19). **f.** Same as e, but for D2 MSNs. CS+ (Pearson correlation coefficient; *P* = 0.0014) and CS- (Pearson correlation coefficient; *P* = 0.81). **g.** D1 MSN with significant change in firing rates to CS-, but not CS+. CS+ (Pearson correlation coefficient; *P* = 0.072) and CS- (Pearson correlation coefficient; *P* = 0.0007). **h.** Same as g, but for D2 MSNs. CS+ (Pearson correlation coefficient; *P* = 0.63) and CS- (Pearson correlation coefficient; *P* = 0.0078). All panels depict data from the first 25 trials of each cue.

**Extended Data Fig. 6.**
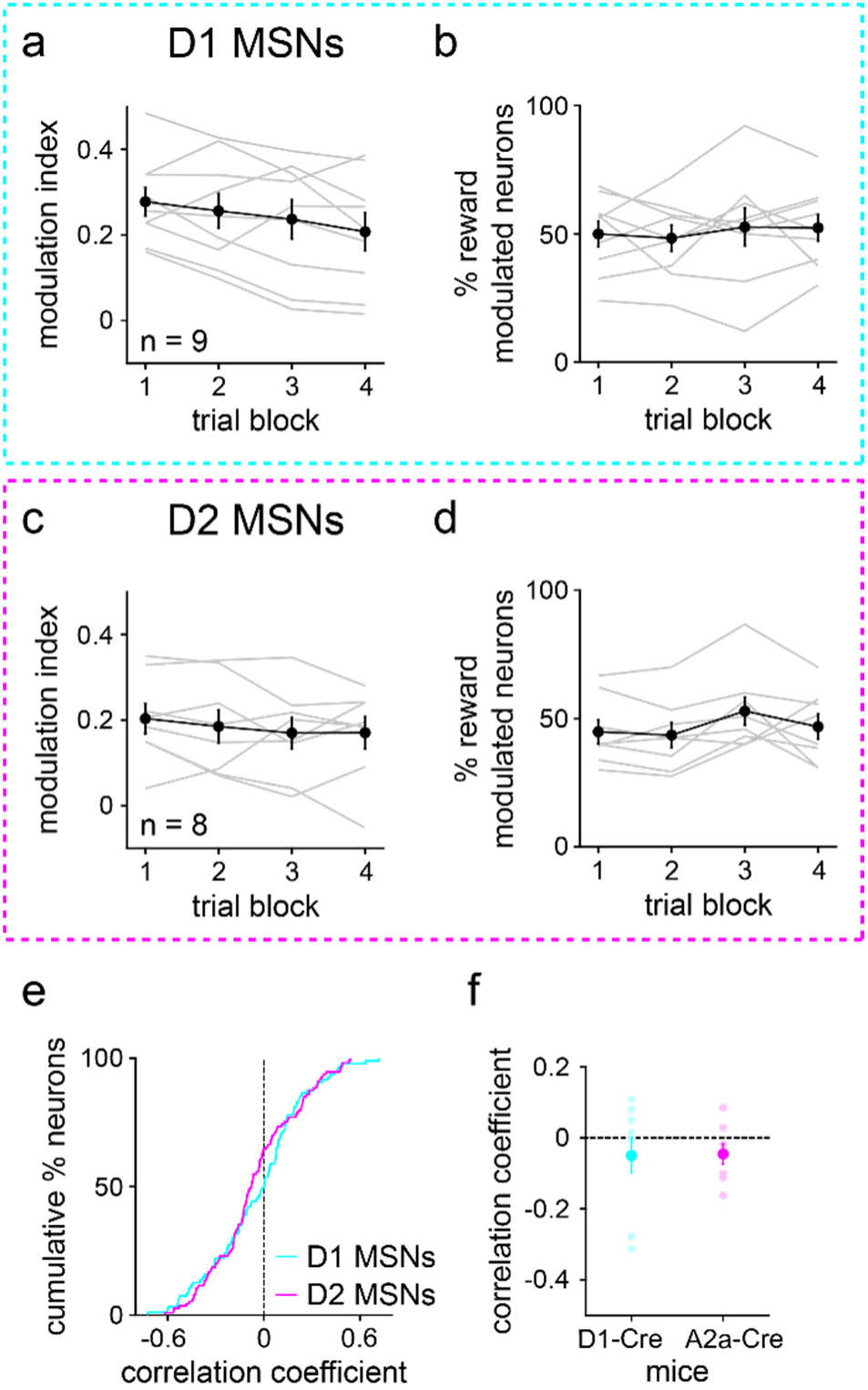
MSNs exhibit fixed average reward activity during learning. **a.** D1 MSN modulation index versus trial block (*n* = 9 D1-Cre mice; one-way RM ANOVA; block effect: *F*_3,24_ = 3, *P* = 0.067). **b.** Percentage of reward modulated D1 MSNs versus trial block (one-way RM ANOVA; block effect: *F*_3,24_ = 0.4, *P* = 0.78). **c.** Same as a, but for D2 MSNs (*n* = 8 A2a-Cre mice; one-way RM ANOVA; block effect: *F*_3,21_ = 0.6, *P* = 0.60). **d.** Same as b, but for D2 MSNs (one-way RM ANOVA; block effect: *F*_3,21_ = 2, *P* = 0.095). **e.** Cumulative percentage of D1 and D2 MSNs versus correlation coefficient between reward firing rate and trial number (first 25 trials) (*n* = 95 D1 and 115 D2 MSNs; two-sample Kolmogorov-Smirnov test; *P* = 0.17). **f.** Correlation coefficient between reward firing rate and trial number (first 25 trials), plotted for each MSN type (*n* = 9 D1-Cre and 8 A2a-Cre mice; two-sample t-test; *P* = 0.94). Dark colors in a-d and f depict mean ± SEM; light colors depict individual animal data.

## ACKNOWLEDGMENTS

This work was supported by National Institutes of Health grants F31MH142117 to T.B.D. and R01DA060229, R01NS125877, R01NS136137 to S.C.M.

## AUTHOR CONTRIBUTIONS

T.B.D. – funding, methodology, data collection, analysis and interpretation, validation, manuscript preparation; J.C. – validation; R.L. – validation; S.C.M. – funding, resources, methodology, analysis and interpretation, manuscript preparation. All authors reviewed the prepared manuscript.

## METHODS

### Animals

Animal procedures were approved by the University of California, Los Angeles, Institutional Animal Care and Use Committee. To record from or optogenetically stimulate genetically defined D1 and D2 MSNs, experiments used male D1- or Adora2a(A2a)-Cre mice or Cre-negative littermates for controls^17^. Mice were maintained as hemizygotes with a C57BL/6J background (strain #000664, The Jackson Laboratory). Mice were maintained on a 12-hour light/dark cycle in temperature- and humidity-controlled housing. Mice were initially group-housed, then singly housed following the first surgery. Mice were aged 8-12 weeks at the time of the first surgery.

### Surgical procedures

Prior to surgery, mice were administered carprofen (5 mg/kg S.C.) and anesthetized with a mixture of 4% isoflurane/oxygen. Anesthesia was maintained using 0.5-1.5% isoflurane. During the initial stereotaxic survival surgery (Model 1900, Kopf instruments), using a pulled glass pipette (3-000-203-G/X, Drummond Scientific) and nanoinjector (Nanoject III, Drummond Scientific), mice were injected with Cre-dependent AAV in ventral striatum (1.1 mm AP, 1.25 mm ML, 200 nL deposited at −4.6 DV and 300 nL deposited at −4.3 mm DV relative to bregma). Optogenetic tagging and perturbation experiments utilized rAAV5/Ef1a-DIO-hChR2(H134R)-eYFP (4-5.3 x 10^12^ vg/mL); control experiments utilized rAAV5/Flex-GFP (8.7 x 10^12^ vg/mL, Gene Therapy Center Vector Core at UNC Chapel Hill). In the same surgery, mice were implanted with head bars for head-fixed behavior and electrophysiology. For optogenetic perturbation experiments, mice were implanted with a ferrule-coupled optical fiber (200 µM, 0.22 numerical aperture, ThorLabs), with the fiber tip positioned at −4.15 mm DV. After surgery, mice were administered daily, post-operative carprofen (5 mg/kg S.C.) for two days, and weight was monitored.

### Behavioral task

Behavioral programs were controlled using custom LabView software (v11.0, National Instruments). Following at least one week of recovery from surgery, mice were food restricted and provided with *ad libitum* access to water. Food restricted mice were maintained at greater than 80% body weight. Mice were gradually habituated to handling, head fixation, and un-cued reward training. During head-fixed reward habitation, a milk reward (5 µL of 10% sweetened condensed milk in water) was delivered via an audible solenoid valve. Mice accessed reward via an infrared lick detection and delivery port positioned in front of the mouth. Mice were habituated for 100 trials per day with progressively increasing inter-trial intervals (15-25 ± 5 s). Mice underwent uncued reward training until they exhibited consistent and robust licking to reward, typically three days. Following habituation, mice were trained on the classically conditioned cue discrimination task. Head-fixed, food-restricted mice were presented with two, randomly interleaved (50% probability) conditioned stimulus olfactory cues predictive of different outcomes^33^. On CS+ trials, an olfactory cue (1 s of 10% isoamyl acetate, diluted 1:3 in air) was consistently followed by milk reward (5 µL) at a fixed temporal delay (2.5 s inter-stimulus interval (ISI)). On CS- trials, a different olfactory cue (1 s of 10% citral, diluted 1:3 in air) was followed by no reward. Mice were conditioned for 200 total trials per day (i.e., 100 trials per cue; 15 ± 5 s inter-trial interval). Anticipatory lick behavior was recorded and quantified as any licks occurring within 100-2500 ms after cue onset (i.e., the ISI, excluding the 100 ms time bin during cue onset). Behavioral data were analyzed using custom MATLAB code (version 2024b, Mathworks).

### Electrophysiology, optogenetic tagging, and spike sorting

Following habituation, and one day prior to the classical conditioning session, a subset of mice underwent a second surgery in preparation for electrophysiological recordings. A 2 mm wide by 1 mm long craniotomy was drilled over the viral injection site, and a ground hole was drilled over the contralateral cerebellum. The surgical site was covered with silicone sealant (Kwik-Cast, World Precision Instruments), and the mouse was allowed to recover overnight.

Electrophysiological recordings were conducted using a custom 128-channel silicon microprobe (model 128D, developed by our group^35^) with an attached optical fiber (200 µm diameter, 0.22 numerical aperture, ThorLabs). The silicon probe consisted of four shanks, spaced 330 µm apart and with electrodes spanning the bottom 775 µm. Electrodes faced toward the optical fiber. The optical fiber tip was centered between the four shanks and positioned approximately 50 µm dorsal and 400 µm caudal to the most dorsal electrodes. This probe enabled recording of a large portion of the ventral striatum. The probe was coated with fluorescent dye (DiI, Thermo Fisher Scientific) for histological analysis and slowly lowered into the ventral striatum. The tip of the silicon probe was positioned in the rostral portion of ventral striatum (1.475 mm AP, 1.25 mm ML, −4.975 mm DV relative to bregma). Thus, the optical fiber tip was nearly centered dorsal to the viral injection site (approximately 1.075 mm AP, 1.25 mm ML, −4.15 mm DV relative to bregma).

After insertion, the probe was left to settle in the brain for 30 minutes, then the mouse was trained on the cue discrimination task with simultaneous monitoring of electrophysiology signals and behavior. Electrophysiological data were collected at a sampling rate of 25 kHz using a multichannel data acquisition unit, RHD recording system (C3316, C3008, E6500, Intan Technologies) and the open-source recording controller software (v3.3.1, Intan Technologies). Following completion of the behavioral task, the opto-tagging protocol was conducted: 200 pulses of 473 nm laser (8 mW power, 10 ms/pulse, 3 s between each pulse) were delivered^38^. The probe was then removed, the mouse was sacrificed using transcardial perfusion, and the brain was dissected for histological confirmation of viral expression and recording location.

Electrophysiological data were analyzed using the open-source programs Kilosort2^49^ and phy^50^. First, data were bandpass filtered (3-pole Butterworth filter, 0.6-7 kHz). Next, spike sorting was performed using Kilosort, which clustered data into putative single units. Phy was used to manually sort through clusters and verify each unit based on the spike waveform and a correlogram with a clear refractory period. Similar to previously established protocols, opto-tagged D1 and D2 MSNs were identified based on (1) activation within less than 6 ms from laser stimulation and (2) greater than 95% correlation between the baseline and laser-evoked spike waveforms. These criteria are at least as stringent as those used in previous striatal opto-tagging studies^22–24^. However, as additional verification, opto-tagging analyses were also performed using more stringent criteria (i.e., activation within less than 3 ms from laser stimulation, see Extended Data Fig. 2). For the main analyses, between 5 and 28 D1 or D2 MSNs were recorded per animal.

### Analysis of neural activity across learning

Neural activity was analyzed using custom MATLAB code. To calculate firing rates, spikes were binned in 100 ms increments using a three-bin sliding window average. Change in firing rates were quantified relative to a baseline period (0-1 s before stimulus onset). To identify changes across learning, the 100-trial training session was parsed into four 25-trial blocks (or 5-trial blocks for the indicated extended data figures). Across-learning means were calculated by averaging results across time bins (100-2500 ms after cue onset for most analysis, or 0-500 ms after reward onset for analysis of reward-evoked activity), then averaging across animals.

### Neural selectivity and modulation indices

The neural selectivity index quantified differences in neural activity evoked by the CS+ relative to the CS- (i.e., positive/negative selectivity indicated greater/lesser change in activity to CS+ relative to CS-). The modulation index quantified differences in neural activity evoked by the CS+ or CS- relative to the preceding baseline period. Both neural selectivity and modulation index were quantified for each time bin by calculating the area under the receiver operating characteristic curve, subtracting 0.5, then multiplying by 2. Significantly selective responses were identified by comparing the observed difference in mean with the resampled difference in mean using 400 permutations of randomly shuffled trials. The neuron was defined as selective or modulated if at least three consecutive time bins exhibited a statistically significant change in firing (greater than 95% difference between the observed and resampled difference in the mean)^51^.

### Optogenetic perturbations

A separate cohort of mice was used for optogenetic perturbation experiments. D1-Cre mice or Cre-negative littermates were unilaterally injected with Cre-dependent virus driving expression of ChR2 or GFP, then implanted with a permanent optical fiber in ventral striatum (see Methods on surgical procedures). After recovery and habituation, mice were trained on the cue discrimination task until there was consistent anticipatory licking to the rewarded cue and suppressed licking to the unrewarded cue, typically two days. On a subsequent day of training, 100% of CS- trials were paired with laser stimulation (2.5 s laser following cue onset; 473 nm, 4 mW power, 25 Hz pulsed). This protocol permitted a within animal comparison of laser versus no laser conditions.

### Histology

Mice were anesthetized with an overdose of isoflurane until areflexic then transcardially perfused with 10 mL of 1x phosphate buffered saline (PBS) followed by 10 mL of 4% paraformaldehyde (PFA) in PBS. Brains were stored in 4% PFA for approximately 16-24 hours, washed three times with PBS, and stored in PBS at 4°C. Brains were sectioned at 100 µM thickness using a vibratome (VT1000S, Leica) and stored in PBS at 4°C. For immunofluorescence staining, brain sections were washed three times in PBS, washed three times in 0.5% Triton X-100 in PBS, and incubated for 30 minutes in 5% normal donkey serum / 0.5% Triton X-100 / PBS. Sections were incubated in primary antibody solution for two to three hours at room temperature then incubated at 4°C overnight. Primary antibody solutions contained 1:1000 dilutions of chicken anti-green fluorescent protein (ab13970, Abcam) and sheep anti-tyrosine hydroxylase (PA1-4679, Thermo Fisher Scientific) in 5% normal donkey serum / 0.5% Triton X-100 / PBS. Following primary antibody incubation, sections were washed three times in PBS then incubated in secondary antibody solution for three hours at room temperature. Secondary antibody solutions contained 1:500 dilutions of donkey anti-chicken AlexaFluor 488 (703-545-155, Jackson ImmunoResearch) and donkey anti-sheep AlexaFluor 647 (713-605-147, Jackson ImmunoResearch). Following secondary antibody incubation, sections were washed three times in PBS, incubated in a 1:5000 dilution of DAPI solution for ten minutes, and washed a final three times in PBS. Sections were mounted on 25 x 75 x 1 mm microscope slides using CC/Mount tissue mounting medium (C9368, Millipore Sigma) and coverslipped with 24 x 60 mm no. 1 cover glass. Sections were imaged using a fluorescence microscope (Axio Examiner.D1, Zeiss) and Zen 2 pro (Zeiss).

## QUANTIFICATION AND STATISTICAL ANALYSIS

Data collection and analyses were conducted without blinding. Statistical analyses were conducted using Prism (v10.6.1, GraphPad Software). Statistical tests, p-values, and sample sizes were provided in figure legends. For nearly all figures, averages and SEMs were calculated on a per animal basis. However, for the extended data figure on cue modulation (Extended Data Fig. 3), data were averaged after pooling neurons across animals due to a limited number of cue-inhibited and non-modulated cells. Changes in behavior/neural activity across trial blocks and number of tagged cells versus maximum latency were tested using one- or two-way independent or repeated measures (RM) analysis of variance (ANOVA). Dunnett’s and Šidák’s multiple comparisons were used with one- and two-way ANOVAs, respectively. Correlations of behavior versus neural activity and firing rate versus trial number were analyzed using the Pearson correlation coefficient. Correlation coefficients between firing rate versus trial number were tested using two-sample Kolmogorov-Smirnov tests (for by cell analyses) and two-sample t-tests (for by animal analyses).

